# Adaptive introgression in subdivided populations leaves distinct genomic footprints under global and local selection: a case study in *Ciona intestinalis*

**DOI:** 10.64898/2026.09.29.754965

**Authors:** Manon Le Goff, Frédérique Cerqueira, Benjamin Penaud, Pierre-Alexandre Gagnaire, Frédérique Viard, Nicolas Bierne

**Author notes:** These authors contributed equally to this work (Frédérique Viard, Nicolas Bierne).

## Abstract

Adaptive introgression, the transfer of beneficial alleles between species through hybridization, is increasingly recognized as an important source of adaptation to novel environments. However, the theory of adaptive introgression has primarily been developed for a single panmictic population, whereas many empirical examples involve spatially subdivided populations. In such systems, a low-frequency adaptive allele may either represent the leading edge of an adaptive travelling wave or occur in a subpopulation where the introgressed allele is locally selected against. We explored how the relationship between adaptive allele frequency and introgressed tract length differs between these two scenarios. During the spread of an adaptive wave, introgressed tracts are progressively eroded by recombination with the genetic background of the receiving species, so that tract length, like allele frequency, reflects the distance traveled by the wave. Under local adaptation, however, allele frequency depends on the local balance between selection and migration, generally breaking this relationship. Simulations predicted a positive correlation between allele frequency and tract length during the spread of an unconditionally advantageous allele, but no correlation under local adaptation. We tested this prediction in a well-characterized case of adaptive introgression in the sea squirt Ciona intestinalis, using haplotagging to reconstruct phased genomes and local ancestry. We found no evidence of a relationship between adaptive allele frequency and introgressed tract length, supporting a model of local adaptation rather than an ongoing travelling wave.

**Author summary:** Hybridization between species can transfer beneficial genetic variants from one species to another, enabling the latter to adapt to a new environment. However, such variants are not necessarily advantageous throughout the recipient species range. In geographically subdivided populations, a low-frequency introgressed allele may either have arrived recently or be selected against locally. Using simulations, we show that combining allele frequency with introgressed tract lengths clarifies this ambiguity, offering a straightforward method to distinguish between globally positive and locally varying selection. When an adaptive allele is beneficial everywhere, it spreads through populations as a travelling wave. Over time, recombination progressively shortens introgressed tracts. Populations reached first have higher frequencies of the adaptive allele sitting on longer introgressed tracts, whereas recently reached populations have lower frequencies and shorter tracts. When selection varies among populations, this relationship disappears because allele frequency reflects local selection rather than the history of the adaptive wave. We tested this prediction in a confirmed case of adaptive introgression in a sea squirt species by characterizing the footprint of introgressed tract length using local ancestry reconstruction. We found no correlation between allele frequency and introgressed tract length, supporting the hypothesis that variation in introgressed allele frequency reflects local selection rather than a traveling adaptive wave.

## Introduction

Adaptive introgression describes the transfer of genetic variants from a donor to a recipient lineage through hybridization, whereby the introgressed alleles are favored by positive selection [1]. Over the past few decades, data provided by genome sequencing has led to a broad consensus regarding the widespread occurrence of hybridization and adaptive introgression in natural ecosystems [2,3]. These evolutionary processes have a significant impact on both population fitness and the genomic landscape of genetic variation. Hybridization has been suggested to play a role in species response to global change, for example by increasing the survival of hybrid individuals through the emergence of new genetic combinations [4], facilitating rapid adaptation to new environments [5], or even by contributing to the management of invasive species by slowing their spread [6]. Adaptive introgression can further enhance adaptive potential by increasing genetic diversity, thereby enabling evolutionary rescue through the incorporation of beneficial alleles [7,8].

In theory, adaptive introgression can be modelled in a manner analogous to a standard selective sweep, except that the beneficial mutation originates by migration from another lineage rather than arising de novo or from standing variation [9,10]. In the selective sweep literature, studies generally focus on how a beneficial allele increases in frequency within a single recipient panmictic population. The frequency of the advantageous allele increases until it reaches fixation, and linked neutral alleles hitchhike along proportionally to their degree of linkage with the sweeping allele [11]. Fewer models also take into account the spread of a favourable allele in subdivided populations. In these models, recombination alters or can even reverse the associations between the favourable allele and linked haplotypes during spread [12]. Furthermore, spread among different demes may be ongoing. The beneficial allele may already be at high frequency in the deme of origin, where it arose through de novo mutation or introgression, at intermediate frequency in nearby demes, and absent or rare in more distant demes. Booker et al. [13] referred to this as the “lag-type” scenario, emphasizing the delay between the initial rise of a beneficial allele in its deme of origin and its subsequent spread and establishment throughout the metapopulation. Finally, the adaptive allele may confer a fitness advantage only in a subset of environments, leading to local rather than global adaptation. Such local adaptations are likely common. For example, an allele conferring resistance to a pesticide may confer a better survival in treated areas only, but be counter-selected in untreated areas [14]. Similarly, one of the most compelling examples of adaptive anthropogenic introgression-that of alleles conferring tolerance to hydrocarbon pollution in the Houston Harbour killifish [7]-fits this scenario of local adaptation as the native allele is found in pristine populations adjacent to the harbour. This scenario implies a cost of resistance/tolerance that reduces the selective value of the allele in untreated/unpoluted areas [15]. In this case, migration–selection balance preserves the adaptive polymorphism, while flanking regions homogenize following the initial local sweep [16,17]. Shortly after the sweep, the covariance of neutral allele frequencies is expected to increase [18], when the chromosomal extent of the differentiation peak between habitats reaches its maximum. Next, the differentiation peak narrows due to gene flow [16,17]. The homogenisation of the sweep shoulders is slowed down by the genetic barrier generated by the selected locus in proportion to the linkage between the adaptive locus and neutral variants [19]. Regardless of whether the allele is unconditionally advantageous or locally adapted, the genomic extent of the hitchhiking footprint of a sweep is reduced by population subdivision compared to the panmictic case [20]. However, spatial variation may differ depending on the speed and distance travelled in the former case and the heterogeneous distribution of suitable habitats in the latter. Furthermore, the allele frequency is no longer informative, since a low allele frequency can be explained by either a recent arrival or a local disadvantage of the allele. The similarities between the two scenarios are particularly expected in cases of recent anthropogenic adaptive introgression, most of which being characterized by introgressed haplotypes that have not yet become established.

Here we first used a forward-in-time simulation approach to show that the two scenarios of global and local introgression differ in their relationship between the length of introgressed tracts and their frequency in different populations. Then we built on these predictions to infer that introgression between the sea squirts *Ciona robusta* and *C. intestinalis* was a case of local adaptation.

We show that the chromosomal signature left by an unconditionally favorable introgressed allele spreading in space (i.e., traveling wave model) differs from the footprint of locally adaptive introgression (i.e., local adaptation model), and this can be used to discriminate between the two hypotheses. We investigated the spatial variation in allele frequency and introgression tract lengths obtained by local ancestry inference to distinguish between two models: (i) traveling wave model : a model of ongoing global introgression in which an unconditionally favourable introgressed allele spreads and has reached different frequencies in different populations, (ii) local adaptation model: a model of local introgression, which assumes that the introgressed allele confers a selective advantage in a specific habitat while being counter-selected in others, with allele frequency variation maintained by the migration– selection balance.

We then applied our framework to a case of adaptive introgression between the ascidians *C. robusta* and *C. intestinalis*. These species diverged approximately 3 million years ago (net molecular divergence : 14.5%) and are located at the far end of the speciation continuum, exhibiting strong reproductive isolation [21], yet recent human-mediated contact has led to the transfer of alleles from the introduced *C. robusta* into native *C. intestinalis* populations [22,23]. Despite the rarity of hybridization in the wild [24], an introgression hotspot has been identified, in some populations of *C. intestinalis,* within a single region of chromosome 5 [23]. Previous works have identified this introgression as adaptive, based on multiple tests of neutrality, and have suggested that positive selection acts on a haplotype that carries a cytochrome P450 2U1 (CYP2U1) gene repeat [25]. Temporal monitoring over approximately 20 generations further revealed remarkable stability in the frequency of the introgressed allele within populations, together with progressive erosion of introgressed ancestry in surrounding genomic regions [26]. While this pattern is consistent with local adaptation, it remains difficult to distinguish from the effects of a recent spread. Here, we inferred local ancestry from phased genomes reconstructed using a linked-read sequencing approach (haplotagging; [27]) for individuals sampled across multiple populations. We then examined the relationship between introgression tract length and allele frequency. The results are consistent with a scenario of local adaptation, providing independent support for conclusions previously drawn from temporal stability.

## Results

### Distinguishing between travelling wave and local adaptation models

We modelled two scenarios of adaptive introgression using forward-in-time simulations in SLiM 5.0 [28]: one in which the introgressed beneficial allele spreads throughout the metapopulation, and another in which it spreads only in demes where it is positively selected, while being maintained at lower frequencies in other demes through migration (Fig 1). In the first scenario (travelling wave model), we consider that the variability in allele frequency between populations is due to a time difference in the arrival of the adaptive allele in the population (i.e., populations with a low allele frequency would have experienced a more recent arrival).

**Fig 1.**
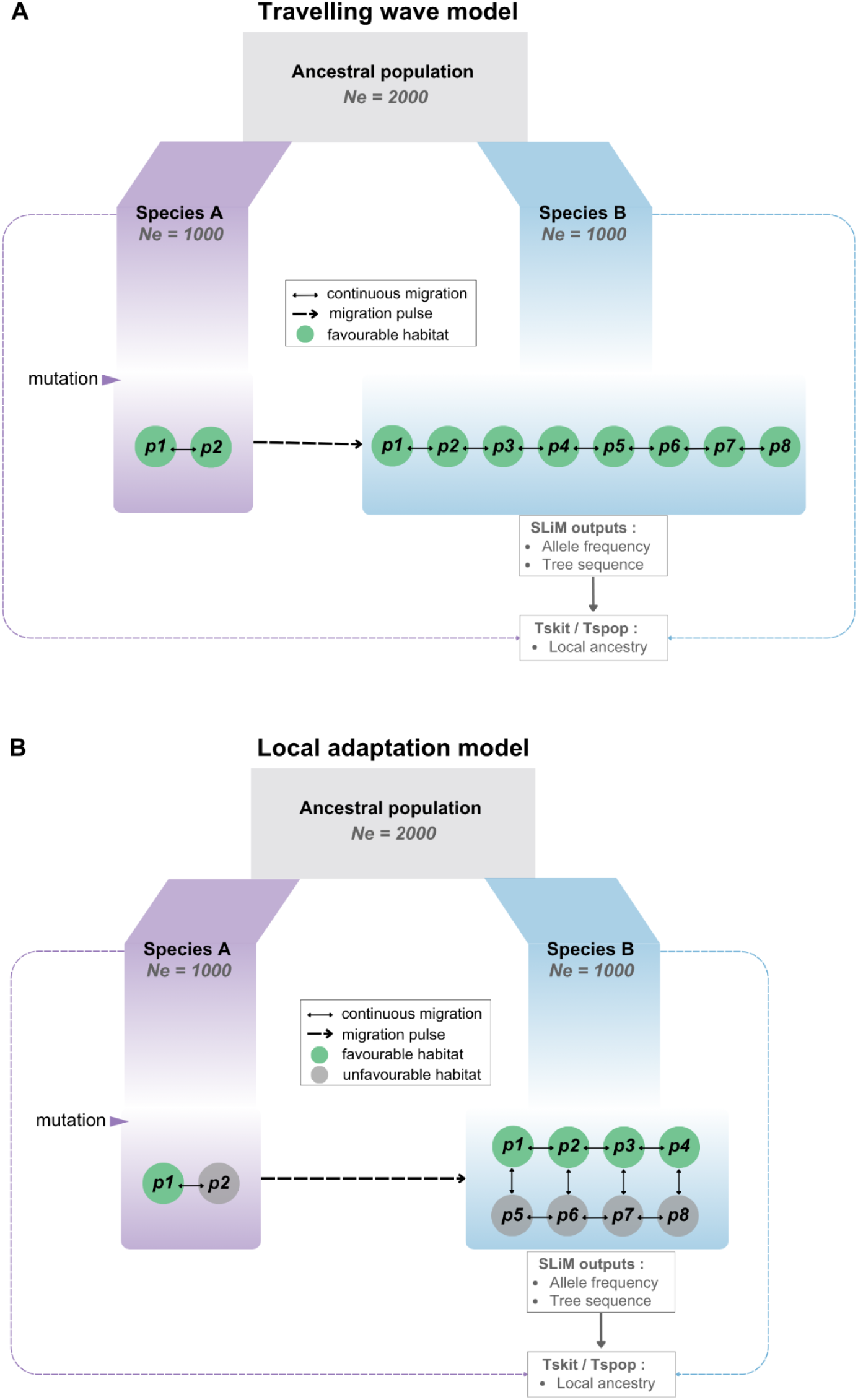
Schematic representation of the two adaptive introgression models simulated with SLIM. (A) travelling wave model, in which the introgressed mutation spreads through the recipient lineage (in blue, hereafter Species B) following a stepping-stone model and is advantageous in all populations. (B) Local adaptation model, in which populations are arrayed along a mosaic habitat and the introgressed mutation is advantageous in half of the populations and disadvantageous in the other half.

In the second scenario (local adaptation model), we consider that the variability in allele frequency is mainly driven by local selection, positive (high frequency) or negative (low frequency). At the end of the simulations, we extracted the local ancestry of the donor species from the recipient species haplotypes to compare the two models. To illustrate the patterns (Figs 2A and 2E), we focused on two recipient populations: one in which the allele reached a high frequency, and one in which it remained at a low frequency. For each population, we represented the two haplotypes of 50 sampled diploid individuals, showing local ancestry with respect to the two parental populations. This initial visualisation already revealed differences between the two models. In populations where the allele remained at a low frequency, introgressed tracts were generally shorter under the travelling wave model than under the local adaptation model.

**Fig 2.**
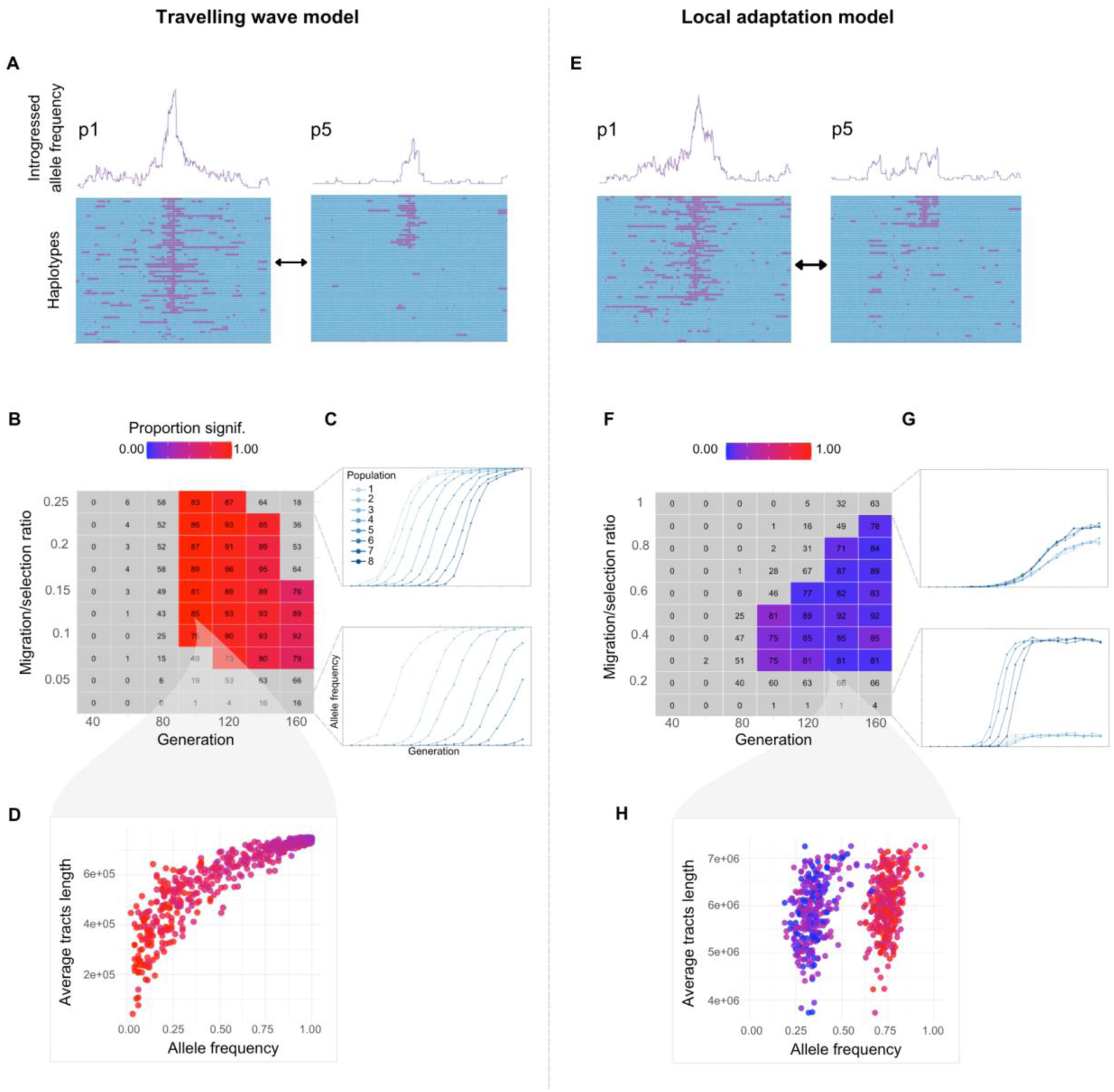
Simulation results for the travelling wave and the local adaptation models. (A, E) Example outcomes showing local ancestry for a single simulation. The purple curve above each graph represents the frequency of introgressed ancestry along the genome. The graphs below show ancestry from the two ancestral lineages, recipient (blue, Species B in Fig 1) and donor (purple, Species A in Fig 1), along the genome for 100 haplotypes (50 diploid individuals) randomly sampled from each population. (B, F) Heatmaps showing the fraction of simulations with a significant correlation between the average length of introgressed tracts and introgressed allele frequency across populations. Each square represents a combination of migration rate, selection coefficient, and generation time. The number inside each square indicates the number of simulations meeting our variability criteria among the 100 simulations performed. (C, G) Examples of the change in introgressed allele frequency over time for the lowest and highest migration rates. Each curve represents a population. (D, H) Examples of the correlation between the average length of introgressed tracts and introgressed allele frequency across populations at generation 90 after introgression, for the 100 simulations.

Based on this observation, we hypothesized that tract length, as a proxy for tract age since admixture, and allele frequency, as a proxy for the time since arrival of the adaptive allele in a deme under the travelling wave model, would be positively correlated under the travelling wave model but not under the local adaptation model. To test for this assumption, we examined the correlation between these two variables across a range of migration/selection ratios. We also monitored how this correlation changed over time (across generations) to detect the effect of the sweep age on it, if any. We observed that a high migration rate reduces the frequency difference between populations and, in the case of the local adaptation model, it also reduces the speed to equilibrium allele frequencies for almost all migration/selection ratios (Fig 2C).

The heatmap (Fig 2B) summarizes the parameter space (migration/selection ratios and time since admixture) in which the allele was present in all populations but not fixed. In practice, we restrict our analyses to parameter combinations for which the adaptive locus is still polymorphic. Under the travelling wave model, the beneficial allele is expected to eventually fix throughout the metapopulation, making the observed polymorphism a transient state. Once fixation is achieved, the selective sweep is complete and there is no longer any local adaptation alternative explanation.

In contrast, under the local adaptation model, polymorphism is maintained at migration– selection equilibrium. The grey squares represent conditions where either the mutation was not yet present in every population, or alleles were fixed for every population (selection dominates migration). Since these conditions depend on the transmission rate of the allele, for the travelling wave model, the squares in the upper right corner (high migration rate and advanced generation) represent cases where all populations were fixed for the mutation. This case does not exist in the local adaptation model, since the allele is maintained at low frequency in populations with a local disadvantage. Conversely, in the bottom left corner, the mutation did not emerge, or at least not in all populations. We are not interested in these cases since we are seeking to study a model that explains allele frequency variability. Each coloured square corresponds to a parameter combination for which this condition was met. The contrast in colour patterns between the two models highlights differences in how the allele behaves under local versus global advantage. The darkest red squares indicate parameter combinations with a high proportion of simulations in which tract length and allele frequency were correlated (Fig 2D). In the local adaptation model, this proportion remained at 0 (or very close to 0) across all tested parameter combinations. In contrast, in the travelling wave model, many parameter combinations resulted in a strong correlation between the two variables, and this occurred in a substantial number of simulations. Similar results were obtained with other connectivity models under the local adaptation model (Suppl. Material, Fig S2)

### Which model better explains introgression patterns in *Ciona* spp.?

Haplotagging data contained sufficient physical phase information for effective haplotype reconstruction and to define introgression tracts. We were able to reconstruct the ancestry of 192 haplotypes despite a substantial reduction in sequencing coverage caused by filtering procedures (see supplementary material). Although this filtering substantially reduced coverage, it resulted in a linked-read dataset of exceptionally high quality. After phasing, we retained 247,378 accurately phased SNPs on chromosome 5. The phased assembly comprised haplotype blocks with a mean length of 54 Kb and an average of 313 SNPs per block. The phase-block N50 reached 7.3 Mb, indicating that a limited number of very long haplotype blocks spanned most of the chromosome and demonstrating the long-range phase resolution. Given that sufficiently long phased haplotypes were obtained despite the relatively low SNP coverage, it was not necessary to include a genotype imputation step, which is an error-prone step in insufficiently sampled non-model species that is often required in haplotagging studies. Based on this phased haplotype dataset, we used Loter [29] to reconstruct the local ancestry of each haplotype. Like many local ancestry softwares, Loter requires parental reference populations to assign each SNP to one of the two parental ancestries. We used three C. robusta individuals as reference, as well as C. intestinalis individuals from the Ettel population, which was the only one showing total absence of introgression in reference-free introgression landscapes obtained with DIEM [30] and polarization analysis.

At the whole-genome level, *C. robusta* ancestry in *C. intestinalis* was almost entirely concentrated on chromosome 5 (Fig 3C), with only very few introgressed tracts elsewhere in the genome that would have allowed us to use the neutral models typically used to estimate the age of admixture.

**Fig 3.**
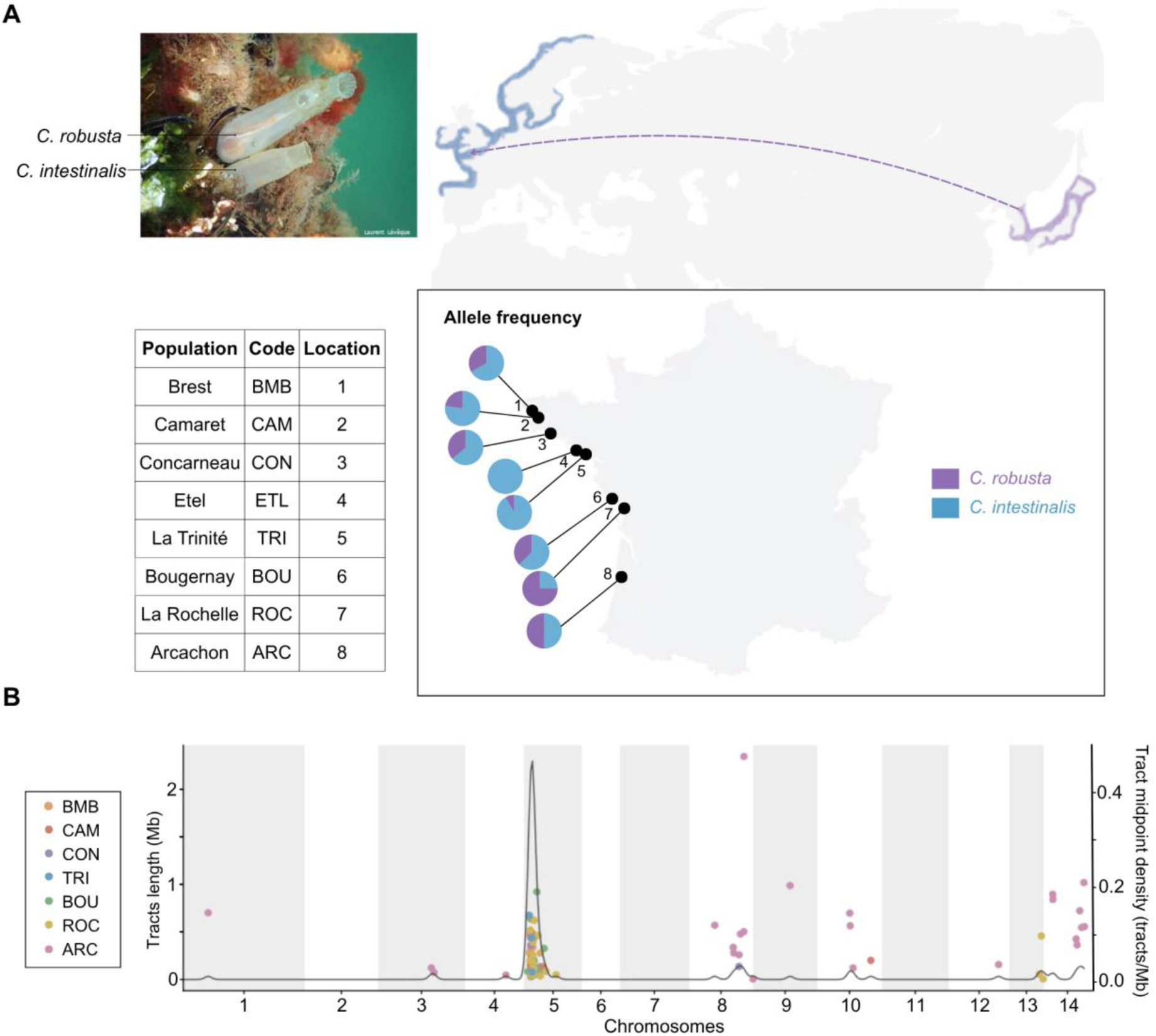
Adaptive introgression on chromosome 5 of *Ciona intestinalis* from the donor species *C. robusta*. (A) *C. robusta* is native to the Northwest Pacific and has been introduced into the native range of the Atlantic species *C. intestinalis*. The photograph shows individuals of both species living at the same location. Pie charts represent the relative proportions of *C. robusta* alleles (purple) and *C. intestinalis* alleles (blue) at the candidate adaptive locus CYP2U1 in each population. Photograph courtesy of Laurent Lévêque. (B) Introgression across the genome. Each point represents an introgressed tract, with tract length on the y-axis and genomic position on the x-axis. Colors indicate populations, and the gray curve represents the density of introgressed tracts across the genome.

We therefore dated the chromosome 5 introgression using DAIM (Deterministic Adaptive Introgression Model) [31], which allowed us to jointly estimate the time since introgression and the strength of selection based on the frequency and length distribution of introgressed tracts. At the metapopulation level, the maximum joint likelihood for *t* and *s* was obtained at 96 generations since introgression, corresponding to approximately 50 years in this species, with an estimated selection coefficient of 0.13. Applying the same approach separately to each locality revealed variation in inferred admixture age among populations, reflecting differences in introgressed tract lengths among localities. This variability between localities also raises questions about the age estimate for the metapopulation. In fact, DAIM assumes a single-population model that does not account for the spatial erosion of introgressed tracts during propagation. Differences in tract length are therefore attributed to differences in the time elapsed since admixture. Although it is possible to impose a common date of admixture on all populations, our theoretical results show that this would not accurately reflect the spatial dynamics of introgression. Thus, 96 generations is our best approximation since it accounts for selection, but a more precise estimate would require an explicit spatial model of adaptive introgression.

Arcachon was the only population that differed in terms of the presence of introgressed tracts outside chromosome 5. Several individuals carried long *C. robusta* tracts elsewhere in the genome, primarily in two regions on chromosomes 8 and 14 (Fig 3C). We considered different possible explanations for this pattern. These tracts might have the same origin as the introgression on chromosome 5. This seems unlikely, since their average length (∼550 Kb) implies a much more recent introgression event than that inferred for chromosome 5. Furthermore, chromosome 5 itself showed no signs of more recent introgression in Arcachon. On the contrary, the introgressed tracts on chromosome 5 were relatively short in Arcachon compared to those observed in other populations. We believe rather that the signals observed on chromosomes 8 and 14 could result from an additional pulse of introgression specific to the Arcachon population. We first assumed this introgression was neutral and calculated the admixture time. Based on the mean length of introgressed segments (545,785 bp) and the genome-wide admixture proportion *v1* = 0.006, excluding chromosome 5, we estimated an admixture time of 19 generations (approximately 10 years) under a neutral model, five times more recent than the event inferred for chromosome 5. However, the concentration of these segments on chromosomes 8 and 14 suggests that the introgressed ancestry is not uniformly conserved across the genome. Heterogeneous selection against introgression may have preferentially eliminated long introgressed segments throughout much of the genome while preserving them in some regions where selection against introgression is weaker, due to higher recombination rates or a lower density of mutations subject to selection. It is also possible that these introgression events, while independent of the one on chromosome 5, are likewise adaptive. For chromosome 8, we did not find a clear introgression peak consistent with the signature of an adaptive introgression event and therefore did not test for potential selection (Fig S6A). For chromosome 14, we identified a modest peak in *C. robusta* ancestry frequency (Fig S6B) and, using DAIM, obtained the joint likelihood surface for the selection coefficient and admixture time and estimated both parameters (Fig S6C). We obtained an estimate of *t* = 29 generations (approximately 15 years) and a selection coefficient of *s* = 0.25. However, these estimates and associated likelihoods should be interpreted with caution, as they may be sensitive to the admixture proportion *v1* = 0.006 used as input in DAIM. In our case, *v1* is difficult to estimate precisely because admixture is not genome-wide but concentrated in a few introgression hotspots. Indeed, an admixture proportion as low as *v1* = 0.05 would be sufficient for neutrality to no longer be rejected (Fig S6B).

We then focused on the first 3 Mb of chromosome 5, as this was the genomic region of interest (i.e., where adaptive introgression took place) (Fig 4).

**Fig 4.**
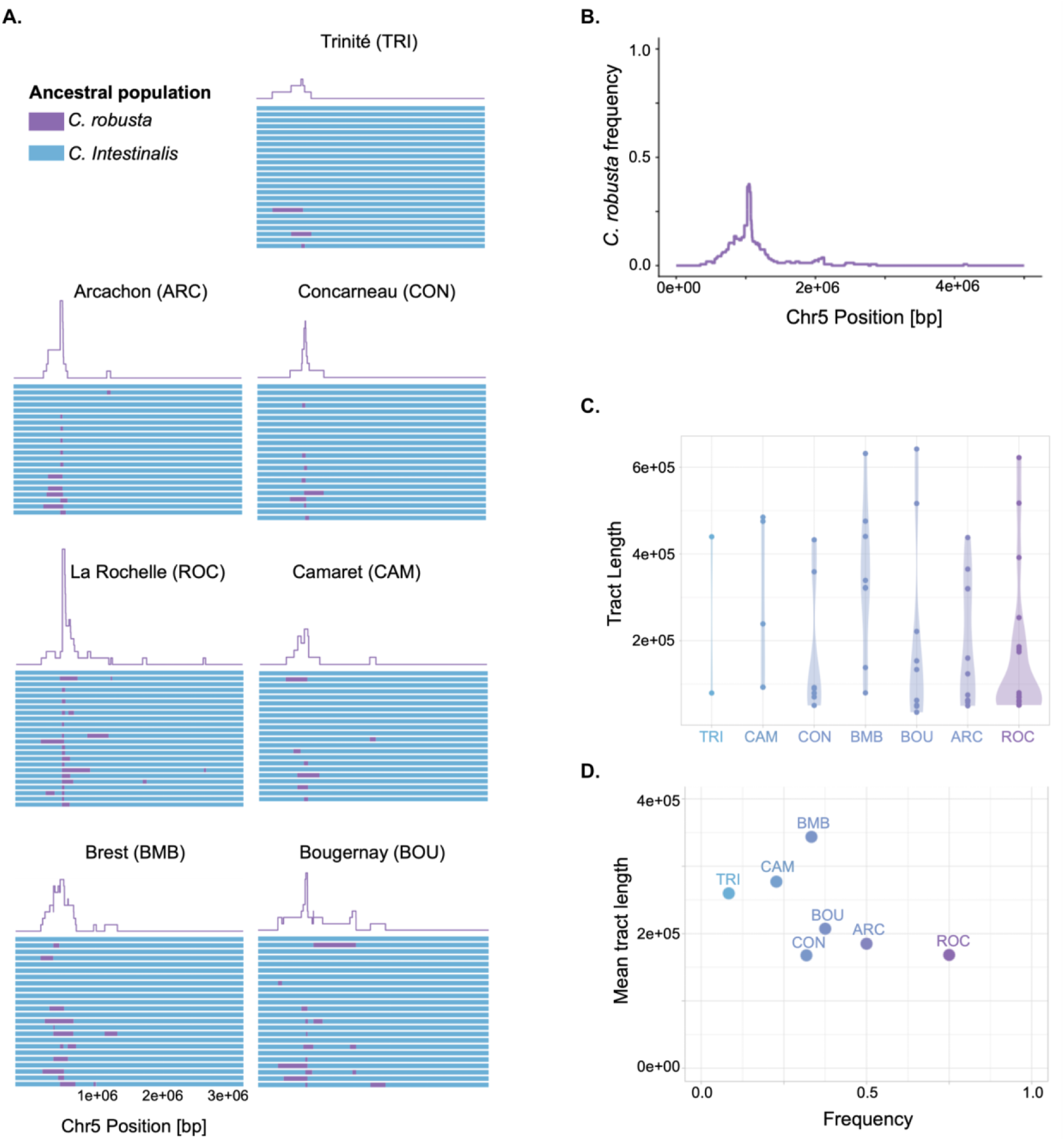
Local ancestry of *Ciona intestinalis* individuals revealed by haplotagging. (A) Each graph shows ancestry from the two ancestral lineages, *C. intestinalis* and *C. robusta*, across the 3-Mb region of interest for 96 *C. intestinalis* individuals studied using haplotagging sequencing, physical phasing and local ancestry inference. The purple curve above each graph represents the frequency of *C. robusta* ancestry along the genome within each population. (B) Frequency of *C. robusta* ancestry along the genome for all the individuals. (C) Length distributions of introgressed tracts carrying the advantageous allele across populations. (D) Average introgressed tract length as a function of allele frequency across populations.

The region identified as having the maximum *C. robusta* ancestry within the *C. intestinalis* species was delimited from position 1,042,729 bp to position 1,055,324 bp. This region corresponds to our expectations since it includes SNP15 (position: 1,055,311 bp), previously used to identify introgressed individuals [26], as well as the gene suspected of being under selection (CYP2U1c, positions from 1,049,202 bp to 1,054,374 bp [32]). Our haplotagging dataset confirmed that the frequency of the introgressed adaptive allele is variable among populations (Fig 3). We subsequently characterised the distribution of mean population tract lengths, as this parameter is expected to correlate with population allele frequency under the travelling wave model. The mean tract lengths (Fig 4C) exhibited minimal difference among populations, whereas the associated distribution displayed greater variability. In light of the results obtained in our simulations, we tested the relationship between introgressed tract length and allele frequency in order to discriminate between the travelling wave and the local adaptation models. To do this, we first grouped haplotypes by population (Fig 4A) and calculated allele frequency and average tract length (Fig 4D). Our data did not reveal any positive correlation between allele frequency and average tract length per population, thus supporting the local adaptation scenario.

## Discussion

Our study shows that population subdivision fundamentally alters the genomic signature of adaptive introgression among demes of a metapopulation. Contrary to the classical expectation derived from single population selective sweep theory, the relationship between allele frequency and introgressed tract length depends on whether selection acts globally or locally across the metapopulation. Under a travelling wave model of a globally advantageous allele, these two quantities remain positively correlated as the adaptive allele spreads across populations. Under local selection, however, this correlation disappears because allele frequency is determined by the local migration–selection balance rather than by the time elapsed since the adaptive allele reached the local deme. These contrasting predictions provide a straightforward conceptual framework to determine the most likely of these two scenarios from genomic patterns of local ancestry in empirical datasets.

### Space, time and introgressed tracts: the genomic signature of adaptive introgression in subdivided populations

The seminal genetic hitchhiking model [11] examines the diversity valley left on a chromosome by the fixation of a beneficial mutation in a single panmictic population. By contrast, under local selection, spatially heterogeneous selection maintains polymorphism at the adaptive locus and a barrier to gene flow in the flanking chromosomal region [33]. In the era of whole-genome sequencing, distinguishing between these two long-term outcomes should, in principle, be straightforward, as one predicts fixation whereas the other predicts persistent polymorphism. However, recent environmental changes, particularly those driven by human activities, create situations in which adaptive alleles may still be spreading toward fixation [13]. When polymorphism is observed, a key question is therefore whether it reflects an ongoing global selective sweep or, alternatively, a locally selected allele approaching a migration–selection equilibrium. This was precisely the question that motivated our work on adaptive introgression in *Ciona intestinalis*: although we found strong evidence for adaptive introgression, the introgressed alleles have not reached fixation [25,26]. Similar situations have been reported in several other examples of anthropogenic adaptive introgression, including the introgression of anticoagulant resistance alleles in mice [34], hydrocarbon tolerance alleles in killifish [7], and insecticide resistance alleles in *Helicoverpa* moths [14]. In these examples, the interpretation of a local adaptation is not much of an issue because the environmental variation and gene function are known and matched. However, the question may arise for cases identified through blind genomic approaches, as was the case for us with *Ciona* spp.

Under the travelling wave model, the adaptive allele progressively spreads across the metapopulation [35]. Its frequency within a deme is therefore expected to reflect the time since the wave reached that deme: high frequencies occur in demes where the allele has had more time to increase under positive selection, whereas low frequencies characterize demes reached most recently. At the same time, non-recombined hitchhiking tracts, which in our case correspond to introgressed tracts, are progressively shortened by recombination as the wave advances across the metapopulation [16,20]. Consequently, the most recently colonized demes should exhibit the adaptive allele at lower frequency and on a more recombined genomic background, resulting in shorter introgressed tracts. This is exactly what we observed in our simulations of the travelling wave model. As noted above, however, this relationship is only transient. It exists only while the adaptive allele is still traveling across the metapopulation and disappears once fixation has been reached in all demes. It also depends on the balance between migration and selection. When migration is sufficiently high and/or selection sufficiently weak, the sweep occurs almost synchronously across demes, leaving little geographic variation and no detectable correlation between allele frequency and tract length. This contrast may be illustrated, for example, by the spread of SARS-CoV-2 adaptive variants. The Alpha (B.1.1.7) variant expanded as a traveling wave across England [36]. By contrast, the Delta (B.1.617.2) variant spread rapidly and almost synchronously across much of the world [37]. This difference probably reflects both increased human mobility following the relaxation of travel restrictions and differences in the selective advantage of the two variants. At any rate, our framework is informative only when adaptive alleles exhibit spatial variation in frequency among demes. Accordingly, we focus on parameter combinations that generate an ongoing travelling wave rather than a nearly synchronous sweep in every demes.

Under the local adaptation model, the frequency of the adaptive allele no longer reflects the time since its arrival in the local deme, but instead the local direction and intensity of selection [38]. The allele reaches higher frequency where it is favored and remains at lower frequency where it is selected against. As a consequence, allele frequency becomes decoupled from introgressed tract length, explaining the absence of correlation in our simulations. Moreover, because the adaptive locus remains polymorphic for long periods, recombination has more time to reshuffle the surrounding genomic background [16,17], allowing introgressed ancestry to persist both over a broader genomic region surrounding the selected locus and across a larger fraction of the metapopulation. Note that spatial reshuffling of introgressed ancestry is also expected under the traveling wave model, but occurs only after the beneficial allele has reached fixation across the metapopulation, once the wave has finished spreading. At that point, a spatial gradient in introgressed tract lengths remains, but subsequent migration progressively homogenizes tract lengths across space, erasing this transient gradient. In any case, this post-fixation process is not relevant to our question. Altogether, our simulations confirm theoretical expectations and show that the joint distribution of allele frequencies and introgressed tract lengths provides a simple way to distinguish transient travelling adaptive waves from migration–selection balance in empirical studies of adaptive introgression.

### Local adaptation in *Ciona* elucidated by combining haplotagging and simulation outcomes

We chose a well-characterized case of adaptive introgression between two highly divergent species, namely the tunicates *C. intestinalis* and *C. robusta*, for which the candidate adaptive region had already been identified [25]. Our objective was to reconstruct local parental ancestry with sufficient resolution to accurately delineate introgressed haplotypes and obtain reliable estimates of introgressed tract length. Achieving this in a diploid species requires phased genomes, which remains difficult to obtain at population scale using conventional short-read sequencing. We therefore used haplotagging, a linked-read sequencing approach that preserves long-range haplotype information [27]. This strategy allowed us to phase hundreds of genomes at moderate cost and reconstruct local ancestry with high confidence. During data processing, we identified a molecular tag-swapping artifact in the original haplotagging protocol [39]. Misassigned reads were exclusively orphan reads lacking companion reads from the same long DNA molecule. Restricting our analyses to linked reads only therefore eliminated this source of error and allowed reliable reconstruction of phased ancestry tracts (see Methods and Supplementary Material). Although this filtering substantially reduced sequencing coverage, coverage remained sufficient because of the relatively small genome size of *Ciona* (170 Mb). More importantly, it produced a refined dataset that was particularly well suited to reconstructing phased haplotypes with minimal phasing errors. Compared with the trio-based (i.e., parents-offspring) phasing strategy used by Fraïsse et al. [25], haplotagging approach provided substantially more accurate ancestry tract reconstruction, even with orphan reads filtering and without the use of imputation.

Dating the introgression of chromosome 5 was challenging because *C. intestinalis* showed virtually no neutral introgression elsewhere in the genome that could provide an independent estimate of the date of admixture. We detected other introgression tracts in the Arcachon population, but we believe that this introgression is independent and does not allow us to date the adaptive introgression of chromosome 5 either. We therefore used DAIM to jointly estimate *t* and *s* based on the selected introgression island itself, rather than relying on a neutral recombination clock as in previous studies [25]. Under the assumptions of a single-population selective sweep model, this analysis dated the introgression on chromosome 5 to approximately 96 generations, corresponding to about 50 years, with an estimated selection coefficient of 0.13. Arcachon was a notable exception, with additional introgressed tracts outside chromosome 5. Their length suggests that they are not remnants of the initial introgression event but result from a more recent pulse of admixture specific to Arcachon, estimated at 19 generations ago under a neutral model. In contrast, chromosome 5 showed no corresponding evidence of recent introgression in this population. These additional tracts may be neutral, occur in genomic segments where selection against introgression is weaker, or result from new adaptive introgression events. Their concentration on chromosomes 8 and 14, particularly chromosome 14, provides some support for the latter hypothesis, although inference of positive selection was hampered by uncertainty in the initial admixture proportions and the evidence remains limited. Future temporal sampling in Arcachon could help distinguish these hypotheses by testing whether *C. robusta* ancestry on chromosomes 8 and 14 progressively declines under selection against introgression or, conversely, develops into distinct introgression peaks under positive selection.

More fundamentally, these inferences of admixture age rely on models of introgression within a single population, without spatial subdivision or environmental heterogeneity. The DAIM method offers a useful way to account for selection when admixture time cannot be estimated from neutrally introgressed segments elsewhere in the genome, but it does not account for the spatial spread and gradual erosion of introgressed tracts among connected populations. Our population-specific estimates illustrate this limitation. Differences in tract length between localities result in differences in the estimated age of admixture, whereas they might instead reflect the spatial dynamics of a single introgression event. The analyses above therefore represent what can currently be inferred using available single-population approaches, rather than a complete reconstruction of the history of adaptive introgression. A fuller understanding of how adaptive introgression spreads and persists within a metapopulation requires explicitly accounting for population structure and spatially heterogeneous selection, which motivated our theoretical analyses.

We therefore turned to the main prediction emerging from our simulations. In line with our expectation under a local adaptation model, we found no relationship between adaptive allele frequency and introgressed tract length. This result is consistent with previous observations suggesting that the adaptive allele does not spread across the species range under uniform positive selection, but is instead maintained by spatially varying selection. Indeed, Touchard et al. [26] showed that the frequency of the adaptive allele at the core of the introgression island remained remarkably stable over nearly 20 generations. The absence of a correlation between allele frequency and tract length therefore provides additional support for the local selection model. Some populations illustrate this particularly well. In La Trinité, for example, the adaptive allele occurs at very low frequency. Under a travelling wave model, such a low frequency would indicate recent arrival of the adaptive allele and should therefore be associated with relatively short introgressed tracts. Based on our simulations, these long tracts at low frequency support the hypothesis of strong counter-selection of the allele in this population. Conversely, in La Rochelle, where the adaptive allele segregates at its highest frequency, introgressed tracts are comparatively short. These results are difficult to reconcile with a simple travelling wave model but arise naturally if allele frequencies primarily reflect local selective conditions rather than the age of introgression.

An additional process that we did not explore here, and that deserves dedicated investigation, may also contribute to the observed pattern: in highly divergent species, adaptive introgression often occurs within a genomic background carrying numerous linked incompatibilities [2]. Selection acting against these linked introgressed variants could progressively eliminate large introgressed haplotypes while favouring recombinant chromosomes that retain the adaptive allele but lose flanking incompatibilities [40]. Such a process predicts that the efficiency of positive selection should increase as introgressed tracts become shorter. In long introgressed haplotypes, positive selection acting on the adaptive locus would be partly offset by negative selection against linked incompatibilities, closely resembling Peck’s "ruby in the rubbish" model [41]. Recombination would progressively release the adaptive allele from this linked incompatibility load by generating shorter introgressed tracts. Interestingly, such a process could explain the weak negative relationship between allele frequency and tract length observed in our empirical data. Explicit modelling of linked selection and the genomic distribution of incompatibilities will be required to evaluate this hypothesis. Regardless of whether this process contributes to the observed pattern, however, our main conclusion remains unchanged, local selection best explains the spatial variation in allele frequency at the CYP2U1 locus in *C. intestinalis*.

More generally, our results suggest that interpreting allele frequencies alone may lead to misleading conclusions about the dynamics of adaptive introgression. Many of the best-known examples of adaptive introgression, such as AHR in *Fundulus grandis* [7], EPAS1 in Tibetans [42], or MUC19 in native americans [43], have primarily been interpreted from differences in allele frequency. In a context where introgression is still ongoing, for example, in the case of recent anthropogenic hybridization, our results show that combining allele frequency with introgressed tract length provides complementary information on the spatial and temporal dynamics of adaptive introgression, revealing patterns that cannot be explained by a travelling adaptive wave and highlighting the importance of explicitly considering local adaptation.

### Incorporating space into the study of adaptive introgression

In this study, we did not attempt to identify the environmental factors underlying the selective advantage of the introgressed allele. Likewise, although our results are most consistent with local selection, we cannot yet identify the environmental conditions under which the allele is selected for and against. The spatial scale of the selective mosaic also remains unknown. Because we sampled only a single microsite within each port, we may have overlooked fine-scale environmental heterogeneity [44], despite growing evidence that strong selective gradients can occur over very short distances in port environments [44,45]. Ports are indeed characterized by steep environmental gradients in temperature, salinity, and heavy metal contamination, particularly between their entrance and inner parts. These gradients result in differences in the composition of the communities established along them (e.g., [45–48]). Regarding specifically *C. intestinalis* in recreational ports, Gauff et al. (2022) showed that individuals transplanted from the entrance to inner locations were disadvantaged, suggesting that the inner-outer pollution gradient may be a selective driver for this species. Characterizing the spatial scale of allele-frequency variation will therefore be an important next step toward identifying the ecological drivers of adaptive introgression in *C. intestinalis*.

Our study also considers a simplified model of population connectivity. In reality, dispersal among ports is largely determined by maritime traffic, with a mixture of short distance and long distance dispersal events [44], rather than by isolation by distance. This dispersal pattern is producing the characteristic mosaic genetic structure (Fig Sup. 4) described in several marine introduced sessile species, including *Ciona intestinalis* [46], *Botryllus schlosseri* [47], and *Didemnum vexillum* [48]. Because these connectivity networks remain poorly characterized, we could not explicitly incorporate them into our simulations. The most realistic scenario is therefore likely to lie between the two extreme models investigated here, combining the propagation of adaptive alleles through shipping networks with locally variable selection after establishment. Encouragingly, our exploration of alternative population structures (stepping-stone, island and mosaic models) indicates that our main conclusions are robust to the underlying connectivity model. Future work integrating realistic connectivity networks with spatially explicit selection should provide a more complete reconstruction of the routes by which adaptive alleles spread through fragmented seascapes.

More broadly, our results suggest that understanding adaptive introgression requires moving beyond the classical view of selective sweeps in a panmictic population toward a spatially explicit theory in which dispersal, recombination, and heterogeneous selection jointly shape genomic patterns of adaptation. Adaptive introgression provides a particularly powerful framework for this purpose because the spread of the adaptive allele is naturally traced by the divergent ancestry of the donor species in the surrounding genomic background. By contrast, inferring the spatial dynamics of a classical sweep arising from a de novo mutation is considerably more challenging because no such straightforward ancestry tracking signal exists and we rely on complex patterns of diversity with varying or even reversing associations [12,13,16,20]. In this respect, anthropogenic introgression under environmental change provides a unique opportunity to observe adaptive alleles spreading through natural populations in real time, offering a rare glimpse of evolution unfolding as a travelling wave across heterogeneous landscapes.

## Material and Methods

### Simulations

We modelled two scenarios of adaptive introgression using forward-in-time simulations in SLiM 5.0 [28]: a travelling wave model in which the introgressed beneficial allele spreads throughout the metapopulation, and a local adaptation model, in which the allele is favored only in a subset of demes (Fig 1). At simulation initialization, we used the initializeTreeSeq() function, which activates tree-sequence recording and enables SLiM’s pedigree-tracking feature to uniquely identify individuals and chromosomes (“haplosomes” in SLiM’s words). Because our downstream analyses relied exclusively on ancestry information recorded in the tree sequence rather than on sequence divergence between species, we did not have to simulate neutral mutation accumulation during divergence. Introgressed tracts were reconstructed directly from the genealogical information recorded by SLiM, with recombination breakpoints defining the boundaries between chromosomal segments inherited from the donor and recipient species. Consequently, neither the absolute divergence time between species nor the accumulation of neutral mutations influences tract reconstruction. We therefore simply set the mutation rate to zero and defined the divergence time between the two parental populations (the donor and recipient species) as 10 generations. We simulated Wright-Fisher populations of *N =* 1000 diploid individuals. We chose a uniform recombination rate equal to 1e-7 and a chromosome size equal to 7,499,939 (which is the size of chromosome 5 of *Ciona intestinalis*, which is our empirical case study - see below). The two parental populations were then split into equally sized demes of 1000 individuals each, two demes for the donor species and eight demes for the receiving species. After 250 generations, we introduced a beneficial mutation into the genome of the first deme of the donor species. By default, SLiM treats the ancestral state as neutral and models the fitness effects of newly arising mutations relative to that state. Here, however, we wished to model a locus experiencing antagonistic selection between two environments. The ancestral allele was advantageous in the original environment (e.g., wild habitat) but disadvantageous in the novel environment (e.g., anthropogenic habitat), whereas the derived allele exhibited the opposite fitness effects. This situation is analogous to a resistance mutation that is beneficial in the presence of a stressor but carries a fitness cost in its absence. This fitness trade-off was implemented in both the travelling wave and the local adaptation scenarios. In the travelling wave model, all demes of the recipient species experienced the novel environment, making the derived allele advantageous throughout the metapopulation and ultimately leading to a global sweep. In the local adaptation scenario, only a subset of populations experienced the novel environment, so that the derived allele was advantageous in some demes but disadvantageous in others. As a result, polymorphism is maintained by migration–selection balance. To simulate such a mutation in SLiM and prevent the initial state from being considered neutral, we used the ‘mutationStack’ function to allow two mutations to occupy the same position (see 14.13 Modeling biallelic loci with a mutation() callback in SLiM manual).

Then, in generation 700, we created a 10-generation contact between the two species in order to simulate introgression within the recipient species. From this contact until the end of the simulation, we tracked the evolution of introgression tracts by recording the frequency of the beneficial mutation within each deme of the recipient species every 10 generations after the initial contact. We also recorded the coalescent histories of our simulations every 20 generations, using the ‘tree sequence’ option in SLiM. We stopped the simulation at generation 900, i.e. 200 generations after contact. To model the first scenario (i.e., travelling wave model), we considered the mutation to be advantageous in all demes of the donor and recipient species, and we used a stepping stone model. For the second scenario (i.e., local adaptation model), we defined the mutation as disadvantageous in half of the demes and advantageous in the other half, and we used a mosaic model to connect the populations (Fig 1). For this second scenario, we also examined other connectivity models, namely an “island” model in which all populations are connected to one another and a “stepping stone” model in which the mutation is advantageous in the first four demes and disadvantageous in the last four. Results of the mosaic model are presented in the main text, whereas results of the other two models are provided in supplementary material.

We analysed the tree sequences of each deme across every generation using tskit [49]. For each deme, we randomly sampled 50 individuals and for each individual in the sample, we identified the corresponding haplosome nodes in the tree sequence. The tree sequence was then simplified to retain only these sampled nodes and the ancestral information necessary to reconstruct their genealogies. Using the tspop package [49], we assigned local ancestry along the chromosome by tracing each genomic segment back to one of the two parental species. This procedure produced ancestry tables containing the genomic coordinates of ancestry tracts (start and end positions) together with their inferred species of origin.

To investigate the conditions under which tract length and allele frequency become associated during adaptive introgression, we explored a range of migration-to-selection (*m*/*s*) ratios. We simulated 10 parameter combinations for each scenario, each replicated 100 times. The selection coefficient was held constant while the migration rate was varied. Different ranges of m/s values were considered for the two scenarios. In the travelling wave model, our objective was to generate substantial variation in allele frequency among demes of the recipient species before the ultimate fixation of the unconditionally favourable allele. We therefore explored *m/s* ratios ranging from 0.025 to 0.25, a range for which an adaptive wave travels through the metapopulation. In the local adaptation model, where polymorphism is maintained by the migration–selection balance, we explored a broader range of *m/s* ratios from 0.1 to 1 in order to assess the effect of migration on the spatial distribution of allele frequencies. For each simulation, we retained only introgressed ancestry tracts carrying the selected allele. The length of each tract was retrieved, and the mean tract length was then estimated for each deme. For every replicate and sampling generation, we quantified the relationship between mean tract length and allele frequency at the selected locus across demes of the receiving species metapopulation. To focus on situations where spatial variation could be meaningfully assessed, we retained only simulations in which the introgressed allele was present in all demes. Demes in which the introgressed allele was close to fixation (frequency > 0.9) were excluded. For each replicate, we calculated Pearson’s correlation coefficient between mean tract length and introgressed allele frequency at the selected locus across demes, together with its associated p-value. Finally, for each combination of generation post-introgression and *m*/*s* ratios, we estimated the proportion of replicates exhibiting a statistically significant correlation (p < 0.05) among the 100 simulation replicates.

### The Ciona spp. case study

#### Reconstruction of phased genomes using haplotagging

In light of the outcomes of our simulation, we examined the case of the adaptive introgression between the ascidians *Ciona robusta* and *C. intestinalis,* by analyzing a dataset produced by haplotagging. Haplotagging is a linked-read sequencing method that relies on tagging long DNA molecules prior to short read sequencing [27]. By assigning a unique barcode to each original DNA molecule, the method preserves long-range haplotypic information, also known as the genomic ‘phase’, even though sequencing is later performed as short reads. Here, we used the protocol proposed by Meier et al. [27].

The technique uses magnetic beads, each of which is covered with barcoded Tn5 transposase. To create unique combinations, the beads were placed in 96 wells with barcode ‘A’ followed by ‘B’, ‘C’ resulting in 885000 unique barcode combinations and one barcode ‘D’ which was used to mark the sample. When high-molecular-weight genomic DNA is added, the long DNA fragments spontaneously wrap around individual beads due to magnetic effect. All the associated DNA fragments will inherit the same molecular barcode during tagmentation: Once bound to the bead, Tn5 simultaneously fragments the DNA and inserts adapters containing both the bead-specific barcode and the sequencing primer sequences. This produces a library of ∼150 bp fragments in which all reads originating from the same long DNA molecule share the same barcode. These barcoded short fragments can then be sequenced using standard Illumina short-read platforms. By reconstructing which reads share the same molecular barcode, it becomes possible to computationally reassemble large haplotype blocks and resolve phase information.

In our case, we used haplotagging to accurately identify the extent of *C. robusta* introgression within the *C. intestinalis* genome at chromosome 5. We used genetic material from 96 individuals collected in 8 locations (Fig 3). Of these, three individuals belonged to *C. robusta* and 93 to *C. intestinalis*. We selected the 96 individuals to obtain a sample that best represented all levels of introgression observed in the study by Touchard *et al.* [26]. In that study, the admixture level of 775 individuals from 22 locations sampled in 2021, in the European contact zone between the two species (i.e., Celtic seas and South European Atlantic Shelf), was analysed using 23 single-nucleotide polymorphisms (SNPs). The frequency of *C. robusta* alleles at SNP15, which is located in the centre of the introgression island, was used to associate each *C. intestinalis* population with a *C. robusta* introgression rate. DNA extraction was performed using the NucleoMag Tissue kit (Macherey-Nagel) or the Monarch kit (New England Biolabs), followed, when necessary, by purification using the SRE XS kit (PacBio) to remove small fragments. The haplotagging library was constructed at the GENSEQ platform (https://genseq.umontpellier.fr/WordPress/) and sequenced at the MGX platform (https://www.biocampus.cnrs.fr/index.php/en/facilities-en/browse-by-facility/platform/357-mgx-en) on a single lane of an Illumina Novaseq S4 flowcell. The protocol was adapted to process 96 samples simultaneously, notably through the use of the Integra Biosciences Mini96 electronic pipette. For the multiplexing of 96 samples, 0.374 ng of DNA was tagged using 1.5 µL of pooled beads (a "D" well). After stopping indexed tagmentation, the samples were pooled by column: 9 pools of 10 samples and 1 pool of 6 samples. An additional 0.8X size selection step was applied to the final pool, yielding a library quantified at 5.8 nM via qPCR (using the NEBNext Library Quant Kit for Illumina).

We processed the raw sequencing reads using the Harpy pipeline (V3) [50], which is specifically designed for processing haplotagging data from demultiplexing and quality control to alignment, variant calling, and phasing. We first demultiplexed individual FASTQ files assigning each barcode combination to a specific BX tag encoding molecule and individual membership. A preliminary quality check verified the data formatting and identified potential library preparation errors. A subsequent quality control step done with fastp (version) [51] removed adapters, duplicates, and low-quality sequences.

All reads that had been trimmed and passed through other QC filters, were aligned using bwa mem (V0.7.17) [52] to the chromosome-scale reference genome of species *C. robusta* (GCA_009617815.1, [53]). We chose the *C. robusta* genome over that of *C. intestinalis* because our study focused on genes introgressed from *C. robusta* into *C. intestinalis* and because it is better assembled (no significant mapping bias was detected by Fraïsse *et al.*[25]). During the mapping we defined the molecules using a parameter of maximum size between mapped reads, which we set at 200,000 bp. At this stage we also added a filter to eliminate unlinked orphan reads (i.e., reads that were found alone on a molecule). Indeed, during data processing we identified an unexpected technical artifact associated with the original haplotagging protocol. The protocol includes a few cycles of PCR amplification performed on pools of 6-10 samples during library preparation. We found that this step introduced molecular tag swapping, causing a small proportion of reads to be assigned to incorrect individuals. Although rare, these erroneously assigned reads are sufficient to bias SNP genotype calls. Fortunately, because of the high combinatorial complexity of the tagging scheme, tag swapping generates almost exclusively orphan reads. We therefore simply retained linked-reads supported by long-range molecular linkage. More details are provided in the supplementary material (Pool-effect).

Then, we performed variant calling using bcftools mpileup and call [54]. The parameter *– populations* was used to compare variants among individuals from the same population, allowing us to differentiate *C. intestinalis* individuals based on their geographical origin. We then filtered the resulting VCF file with bcftools to remove low-quality and incorrect variant sites using the parameters *MAF>=0.05* and *F_MISSING<=0.3* and continue phasing only on chromosome 5, where introgression occurs.

The next step of the standard phased data reconstruction pipeline typically involves a step to impute missing genotypes using statistical inference (usually with STITCH [55] in the case of haplotagging data). After testing various imputation parameters, we decided not to use the imputed data, which resulted in the removal of certain ancestry tracts due to an excessive number of missing genotypes.

Finally, we reconstructed the haplotype phase with HapCUT2 (V1.3.4) [56]. This initial phasing uses physical phase information on molecules tagged during haplotagging. Then we used shapeit4 (V4.2.2) [57], to perform chromosome-wide statistical phasing based population-scale linkage disequilibrium information.

We used Loter 1.0.1 [29] to reconstruct the local ancestry of each haplotype. First, we defined the reference populations for both species. The *C. robusta* reference population consisted of three individuals of the species. For the *C. intestinalis* reference population, we selected all individuals from the Etel population, which was the only one showing no signs of introgression, based on the frequency of the allele under sélection. Then, we used the phased vcf file from shapeit5. Loter allowed us to calculate, on the basis of inference, the association of each SNP with a reference source population and to produce a matrix indicating the origin of each haplotype. For our data, the three available *C. robusta* individuals were used as a reference for this species and all individuals from the Etel population for the species *C. intestinalis*. Across the entire genome, we identified the tracts with *C. robusta* ancestry and calculated their lengths.

### Dating adaptive introgression under a single-population model

In a standard admixture analysis, genome-wide neutral introgression can be used to estimate the initial proportion of admixture, *v1*, and the time since admixture, *t.* These parameters can then be fixed when analyzing individual genomic regions for adaptive introgression, for example using ancestry-hmm-s [58], by comparing a neutral model to a model that includes selection and estimating the selection coefficient, *s*. This approach could not be applied to *C. intestinalis* because, with the exception of the Arcachon population, the introgressed tracts were almost exclusively confined to the adaptive introgression island located on chromosome 5, leaving no genome-wide neutral background from which *v1* and *t* could have been estimated independently. Previous estimates of the age of this introgression, based on a neutral recombination clock [25], are therefore potentially problematic, as the introgression on chromosome 5 is subject to selection.

Therefore, we used DAIM [31] to jointly estimate the time since admixture, *t*, and the selection coefficient, *s*, from the frequency and length distribution of introgressed tracts on chromosome 5. For a fixed initial admixture proportion *v1*, we used tract_length.py to generate predictions over a two-dimensional grid of *t* and *s* values. For each combination of parameters, DAIM predicts both the expected frequency of the introgressed allele and the expected distribution of introgressed tract lengths after recombination and selection. Predictions were interpolated between grid points and compared with the observed allele frequency and tract-length distribution to obtain a joint likelihood surface for *t* and *s*. The maximum of this surface was used to obtain joint estimates of the age of introgression and the strength of selection. Because *v1* could not be estimated from a neutral genomic background, we fixed it to a low value consistent with rare initial introgression and repeated the analysis over several low values of *v1* to assess the sensitivity of the estimates to this assumption. Estimates of *t* and *s* for the adaptive introgression on chromosome 5 were robust to the values tested. Analyses were performed both at the metapopulation level, pooling all individuals as a single population, and separately for each locality.

Arcachon was also treated separately because it was the only population in which we detected substantial introgressed tracts outside the chromosome 5 island. These tracts could reflect proximity to the population in which the original contact with *C. robusta* occurred, additional and possibly more recent episodes of adaptive introgression at other genomic locations, or a more recent pulse of neutral admixture. We therefore estimated the age of this additional admixture under a neutral pulse-admixture model using the genome-wide introgressed tracts outside the chromosome 5 island. Under this model, the expected mean tract length is *Lˉ*=(1-*v1*)*r*(*t-*1]-1, where *r* is the recombination rate, *v1* is the admixture proportion and *t* is the number of generations since admixture [59]. Finally, because chromosome 14 contained particularly strong local concentrations of introgressed ancestry in the Arcachon population, we tested this chromosome for evidence of adaptive introgression by examining *C. robusta* ancestry frequency along the chromosome and using DAIM. In contrast to chromosome 5, estimates for chromosome 14 were more sensitive to the assumed admixture proportion. We first used the genome-wide admixture proportion *v1* estimated across all chromosomes except chromosome 5, and then determined the value of *v1* at which the neutral model was no longer rejected.

### Comparing predictions of the travelling wave and local adaptation models in subdivided populations

We applied the same approach as in the simulations to examine the relationship between the frequency of the introgressed allele and the length of the tracts carrying it. Following previous studies [25,26], we used the frequency of the tracts carrying the advantageous allele that we considered being the CYP2U1 locus. To compute the frequency, we calculated the proportion of introgressed haplotypes carrying it per population. We also measured the length of the *C. robusta* ancestry tracts passing through the CYP2U1 locus, that is, the tracts carrying the selected allele. For each population, we computed the average length of these tracts, yielding one data point per population, as in our simulations.

## Data Availability

The raw data from haplotagging sequencing, are currently being deposited in the ENA (European Nucleotide Archive) database. All genetic data analysis scripts, as well as the SLIM simulation scripts, are available on GitHub: https://github.com/ManonLeGoff/IntrogressionCiona

## Author contributions

Manon Le Goff (Investigation, Formal analysis, Writing–original draft, Writing–review & editing), Frédérique Cerqueira (Investigation, Writing–review & editing), Benjamin Penaud (Software, Writing–review & editing), Pierre-Alexandre Gagnaire (Methodology, Investigation, Writing–review & editing), Frédérique Viard (Conceptualization, Methodology, Investigation, Funding acquisition, Supervision, Writing–original draft, Writing–review & editing), Nicolas Bierne (Conceptualization, Methodology, Investigation, Formal analysis, Funding acquisition, Supervision, Writing–original draft, Writing–review & editing).

## Acknowledgments

The authors are grateful to the GENSEQ platform of the Research Support Unit MEEB for access to equipment and for the generation of the Haplotagging data. We are particularly grateful to Erick Desmarais and Anaïs Bordes, who contributed to the implementation of the Haplotagging protocol at the GENSEQ platform, making it available to the biodiversity and evolution research community in Montpellier. We thank the Montpellier Bioinformatics Biodiversity platform at ISEM for its support with bioinformatics analyses.

## SUPPLEMENTARY MATERIAL

### Pool-effect

During the initial PCA performed to characterise neutral genetic diversity in *C. intestinalis*, a few individuals appeared displaced towards the *C. robusta* cluster, as would be expected for hybrid individuals (Fig S1 “PCA before filtering”). This pattern was unexpected because these individuals originated from populations where hybridization is unlikely and were not detected as hybrids with a KASP SNP assay [26]. In addition, neither biological nor sampling explanations could account for their intermediate position in the PCA. Our investigations showed that this pattern resulted from a technical artifact associated with a pooled PCR amplification step used during library preparation. The pool effect is clearly illustrated in the PCA (Fig S1), where individuals cluster according to their PCR pool (colour of the points) rather than their population of origin (shape of the points).

Following the protocol of [39] a step of PCR amplification of libraries was performed on pools of eight or ten samples (11 cycles recommended in the protocol of Kucka and Chan 2024, 10 cycles in Meier *et al.* 2021 [27], we did 8 cycles). We subsequently found that this procedure resulted in a low level of read swapping among individuals within the same pool, artificially increasing their genetic similarity. We suspect that this artifact originates from residual free transposase complexes remaining after tagmentation. Because these complexes carry the Illumina adapters, they may occasionally promote incorrect assignment of reads during PCR amplification due to tag swapping. Most misassigned reads should thus be associated with unique molecular identifiers (MI tags), whereas genuine tagmented fragments are expected to generate multiple reads sharing the same molecular identifier. We therefore refer to these reads as *unliked orphan reads*.

To remove this source of technical noise, we modified the align_bwa.smk workflow of Harpy (version 3). After mapping and standard Harpy filtering steps, reads were classified according to the number of reads associated with their molecular identifier (MI tag), using the information contained in the bxstats file. Reads belonging to molecules represented by a single read were written to a separate BAM file and excluded from all subsequent analyses, whereas only molecules represented by at least two reads were retained.

This filtering reduced the average sequencing depth from approximately 10X to 2X, corresponding to an effective fivefold loss of coverage. However, it completely eliminated the pool effect (see Fig S1 “PCA after filtering”). After filtering, the three displaced individuals returned to their expected position within their population of origin, and the clustering of individuals by PCR pool disappeared. Instead, individuals clustered according to locality, as expected from the underlying population structure (Fig S1). These results indicate that removing singleton molecules efficiently eliminated the PCR-induced artefact while preserving the genuine biological signal.

**Fig S1.**
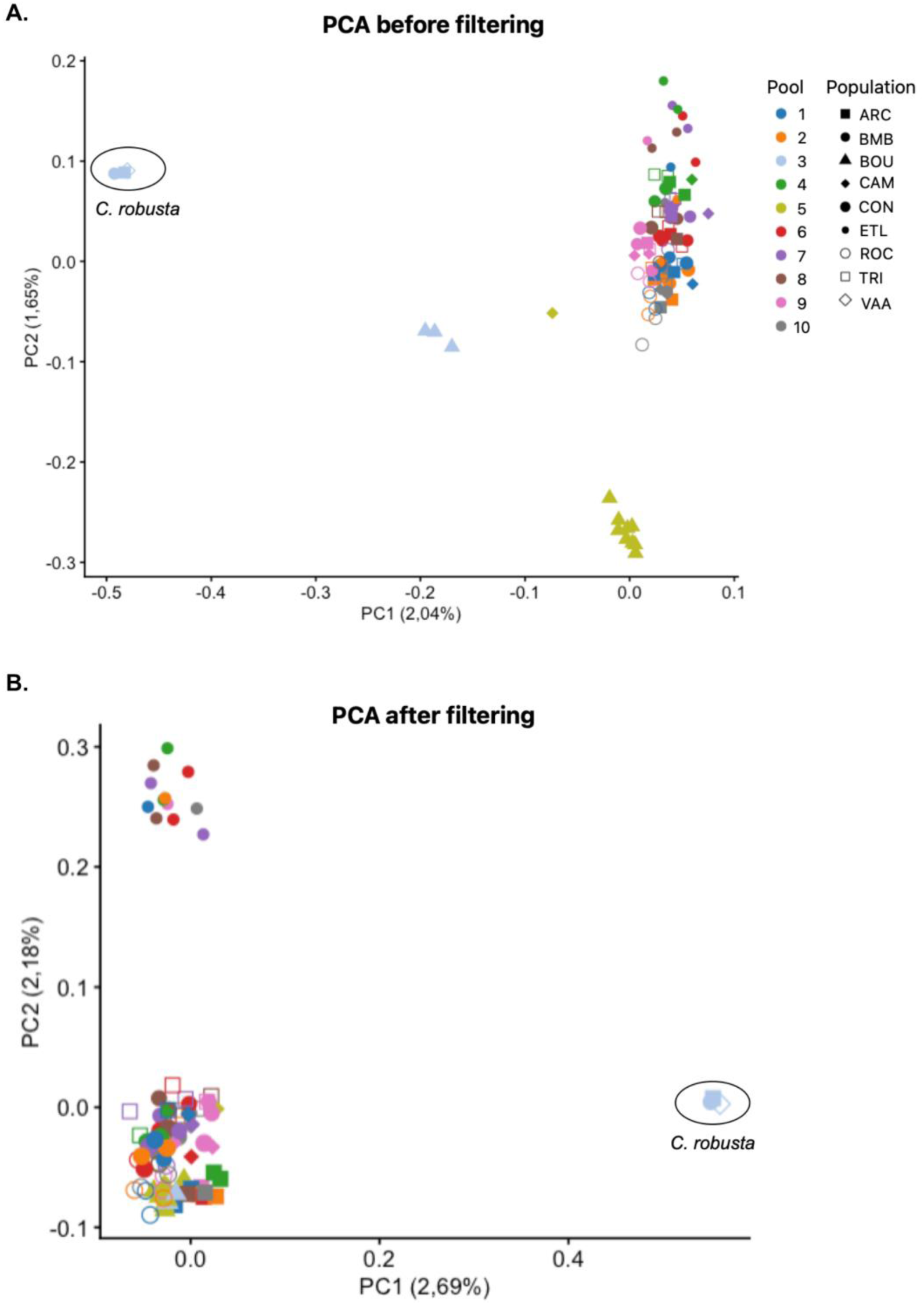
PCA before and after singletons (i.e., orphan reads) filtering. (A) PCA before filtering. (B) PCA after filtering. Each color represents a pool of samples in the 8 cycles PCR amplification. The shapes represent the locality where the individual was sampled.

**Fig S2.**
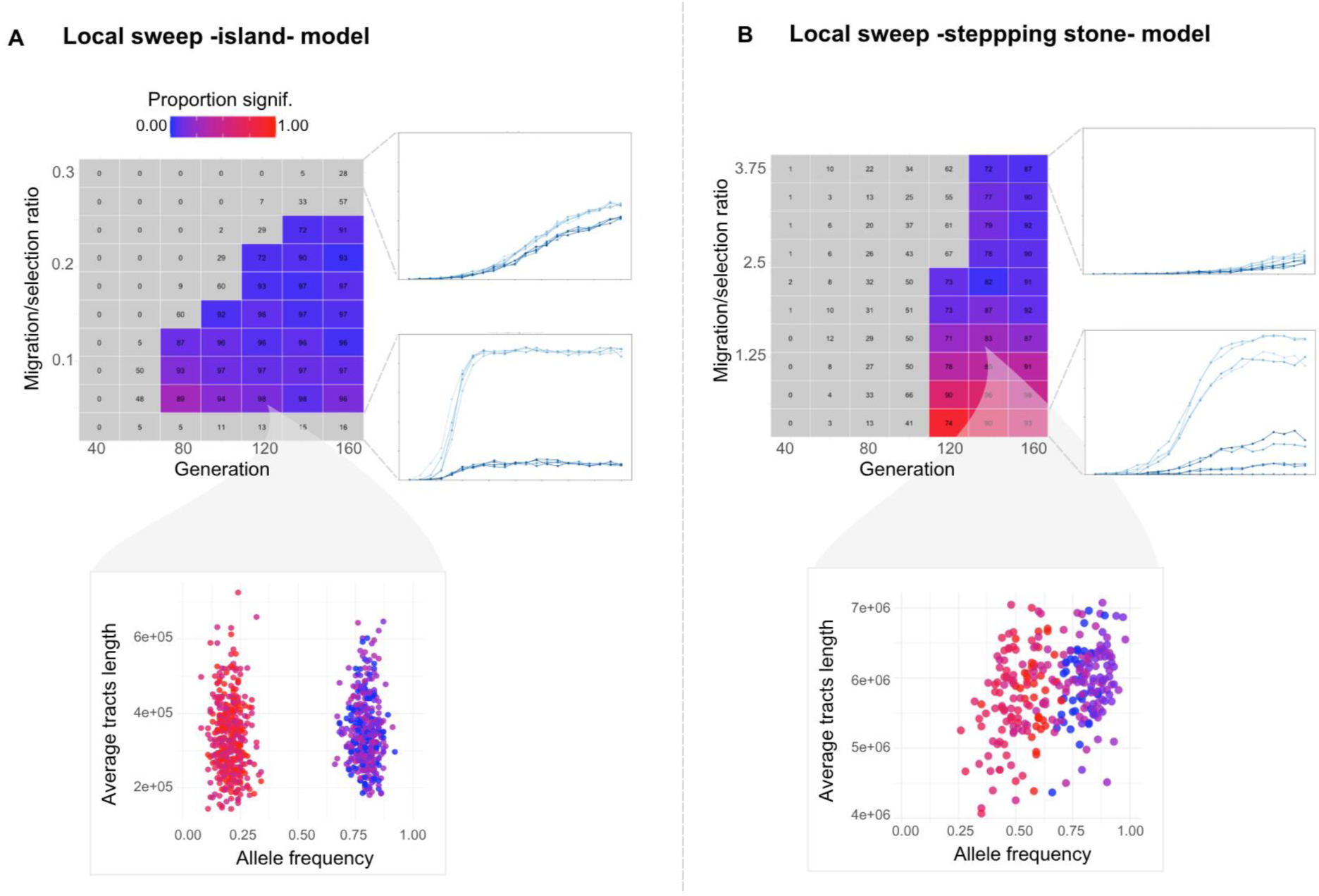
Alternative connectivity models for the local adaptation. Results presented in the main text are based on the mosaic model (Fig. 1). Two additional connectivity models were examined for the local sweep. (A) Island model, in which all populations are interconnected and the introgressed allele is disadvantageous in half of them. (B) Stepping-stone cline model, in which the introgressed allele is advantageous in four consecutive demes and counter-selected in the next four demes. Results were similar to those obtained with the mosaic model, except that positive correlations could arise in the stepping-stone cline model at low migration rates. This results from a global sweep-like effect within the first habitat, followed by a gradient of introgressed ancestry in the second habitat, which can generate a positive correlation between tract length and allele frequency. As migration increases, differences in tract erosion among populations become smaller and this effect disappears.

**Fig S3.**
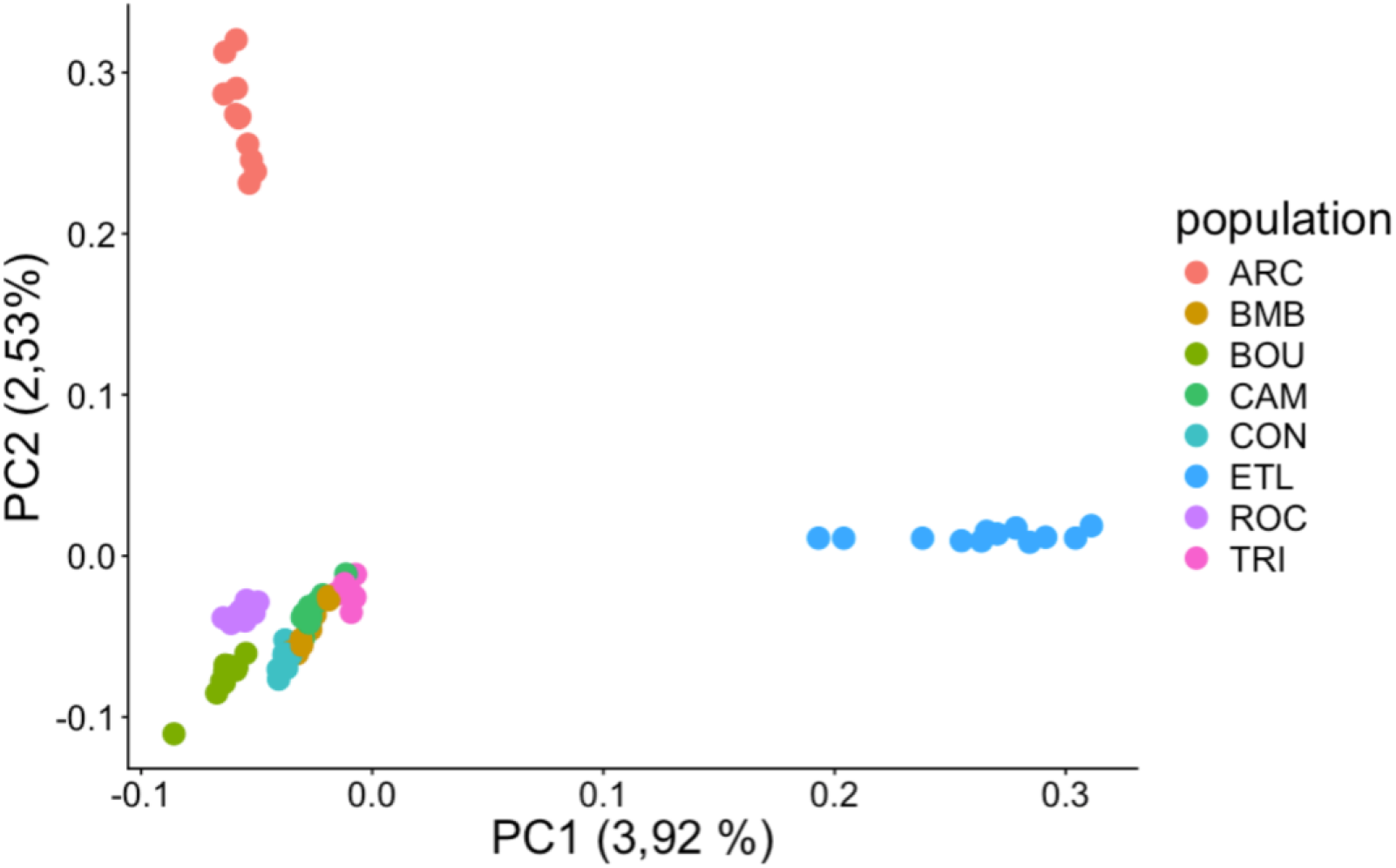
Neutral structure of *Ciona intestinalis.* This PCA was performed on the final dataset after removing chromosome 5 to characterize population structure independently of the major adaptive introgression signal. *C. robusta* individuals were excluded to prevent divergence between the two species from dominating the analysis. The PCA illustrates the mosaic genetic structure described in the main text. Populations do not follow a geographical gradient and are generally weakly differentiated, with the exception of Etel and Arcachon, which stand apart from the other populations.

**Fig S4.**
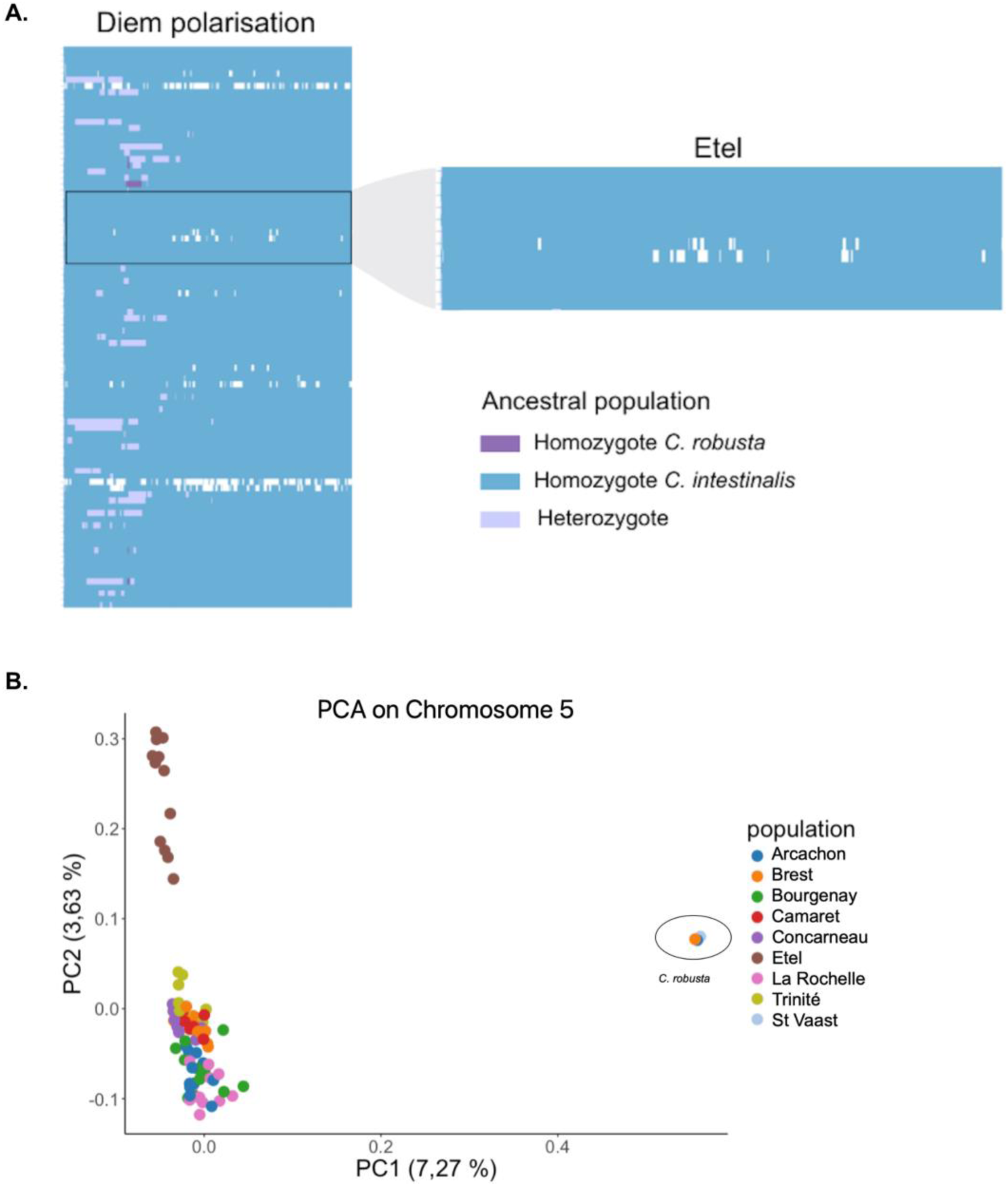
Diem polarisation and PCA on chromosome 5. These two Figs illustrate the choice of using Etel as the reference population for *C. intestinalis*. A) Diem [30] was initially used because it infers ancestry from unphased diploid data without requiring a reference parental population, allowing us to confirm the absence of detectable introgression in the Etel population. However, Diem requires prior screening for ancestry-informative SNPs and is not optimal for handling missing data. For local ancestry inference, we therefore preferred LOTER [29], which infers haploid local ancestry from phased genomes using parental reference populations. LOTER proved more robust for reconstructing introgressed tracts, with fewer interruptions within otherwise continuous tracts. B) PCA on chromosome 5 also shows that the Etel population stands apart and is more distant from *C. robusta* than the other populations.

**Fig S5.**
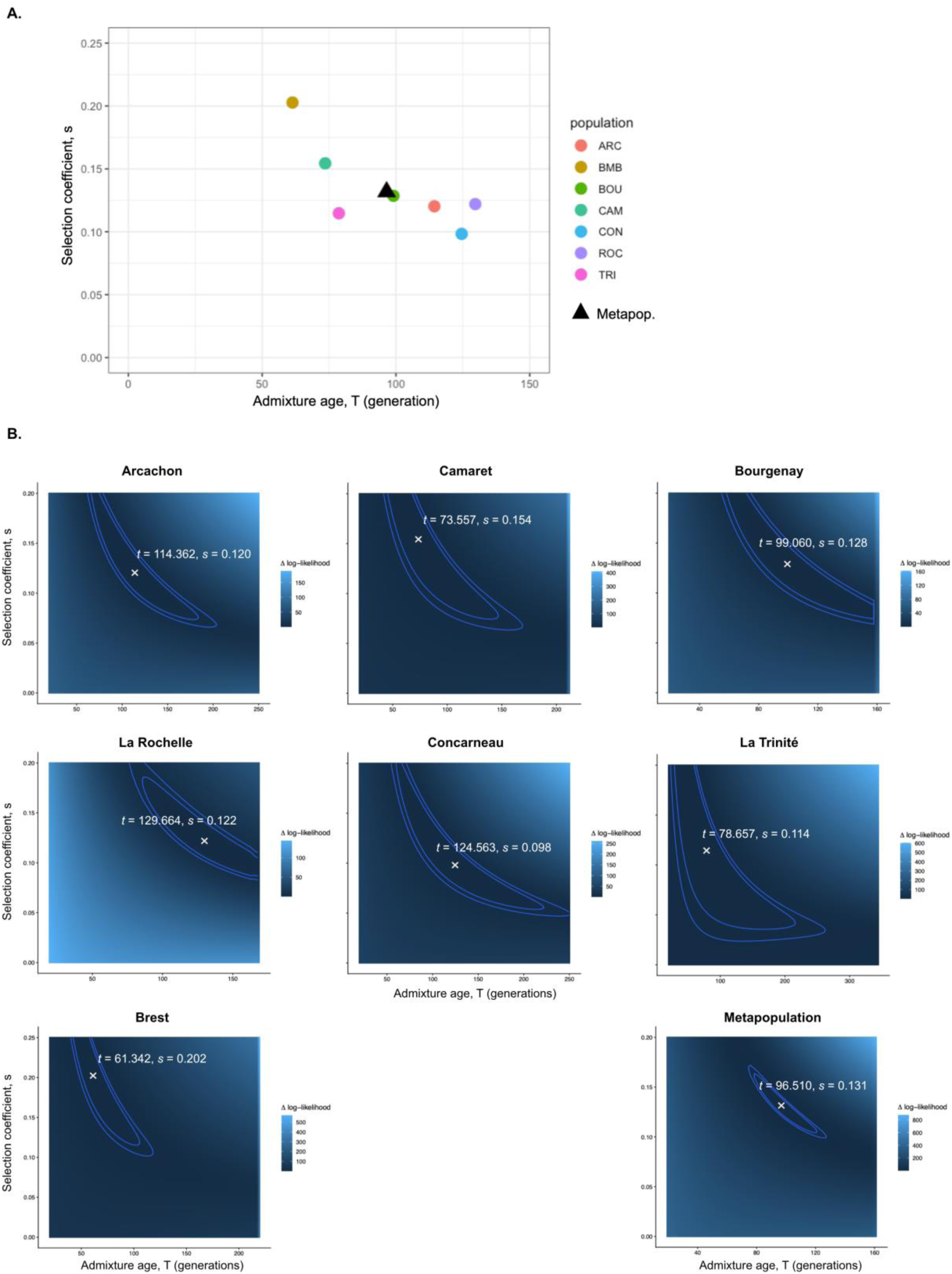
Selection coefficient and introgression time inferred with DAIM. [31]. Each point corresponds to the estimates of the selection coefficient and introgression time for a population, and the triangle corresponds to the estimates obtained at the metapopulation level, with all individuals pooled and considered as a single population. (B) Likelihood surfaces for the selection coefficient and introgression time for each population. The last panel (“all”) corresponds to the metapopulation.

**Fig S6.**
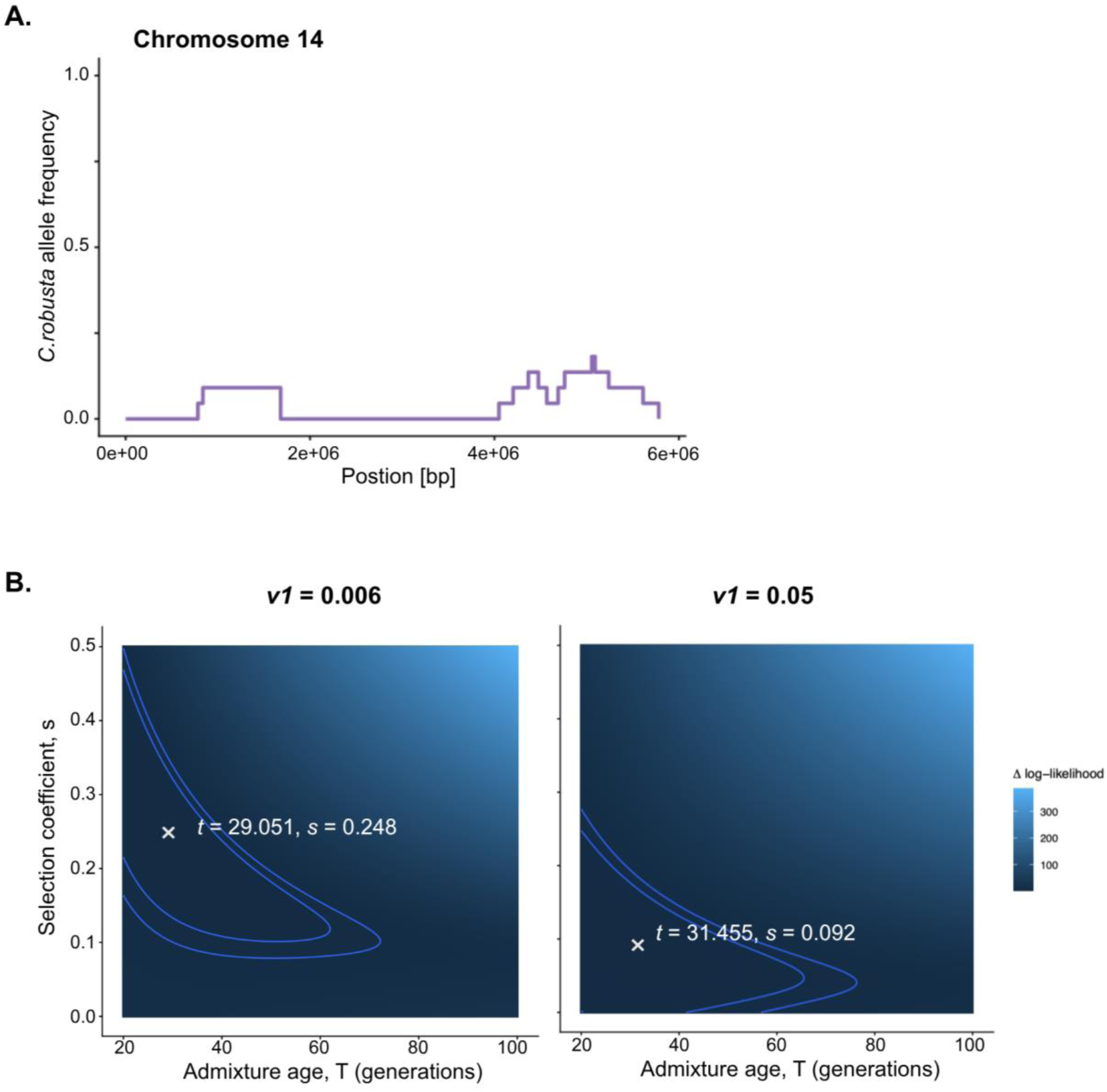
Introgression of chromosomes 14 into the Arcachon population. (A) *C. robusta* ancestry frequency along chromosome 14. (B) Likelihood surface for the selection coefficient and admixture time inferred with DAIM from the introgression peak observed on chromosome 14 with *v1* (genome-wide admixture proportion) equal to 0.006 or 0.05.

